# Dynamic combination of sensory and reward information under time pressure

**DOI:** 10.1101/253641

**Authors:** Shiva Farashahi, Chih-Chung Ting, Chang-Hao Kao, Shih-Wei Wu, Alireza Soltani

## Abstract

When making choices, collecting more information is beneficial but comes at the cost of sacrificing time that could be allocated to making other potentially rewarding decisions. To investigate how the brain balances these costs and benefits, we conducted a series of novel experiments in humans and simulated various computational models. Under six levels of time pressure, subjects made decisions either by integrating sensory information over time or by dynamically combining sensory and reward information over time. We found that during sensory integration, time pressure reduced performance as the deadline approached, and choice was more strongly influenced by the most recent sensory evidence. By fitting performance and reaction time with various models we found that our experimental results are more compatible with leaky integration of sensory information with an urgency signal or a decision process based on stochastic transitions between discrete states modulated by an urgency signal. When combining sensory and reward information, subjects spent less time on integration than optimally prescribed when reward decreased slowly over time, and the most recent evidence did not have the maximal influence on choice. The suboptimal pattern of reaction time was partially mitigated in an equivalent control experiment in which sensory integration over time was not required, indicating that the suboptimal response time was influenced by the perception of imperfect sensory integration. Meanwhile, during combination of sensory and reward information, performance did not drop as the deadline approached, and response time was not different between correct and incorrect trials. These results indicate a decision process different from what is involved in the integration of sensory information over time. Together, our results not only reveal limitations in sensory integration over time but also illustrate how these limitations influence dynamic combination of sensory and reward information.

## Introduction

For many decisions that we face in daily life, collecting more information about the options or actions under consideration is instrumental to making better choices. However, in addition to the mental and physical efforts required to gather and integrate information over time, collecting more information costs time that could be used to make other choices with potentially rewarding outcomes (opportunity cost). Therefore, there is a tradeoff between integrating more information for improved accuracy and faster response time based on information already collected, often referred to as speed-accuracy tradeoff (SAT). The SAT has been the focus of intense experimental and theoretical work over past decades (Bogacz, Wagenmakers, et al., 2010; Chittka, Skorupski, et al., 2009; Miller & Katz, 2013), revealing mental processes (Bogacz, Wagenmakers, et al., 2010; Ratcliff & Smith, 2004; Usher & McClelland, 2001) and neural mechanisms underlying decision-making (Gold & Shadlen, 2007; Huk & Shadlen, 2005; Shadlen & Newsome, 2001).

An important question raised by SAT investigation is whether decision making involves perfect or leaky integration of sensory information over time (Carland, Marcos, et al., 2016; Cisek, Puskas, et al., 2009; Kiani, Hanks, et al., 2008; Stanford, Shankar, et al., 2010; Thura, Beauregard-Racine, et al., 2012; Thura, Cos, et al., 2014; Tsetsos, Usher, et al., 2011) or alternatively relies on stochastic transitions between discrete attractor states (Miller & Katz, 2013). Until recently, a class of models based on perfect integration—namely drift diffusion models (DDMs)—has dominated the field for a few reasons. First, it has long been shown that perfect integration is optimal in the absence of biological constraints such as internal noise (Bogacz, Wagenmakers, et al., 2010). Second, the basic DDM is easy to apply and produces analytical formulae that can be fit to behavioral data (Ratcliff & McKoon, 2008). Third, trial-averaged neural activity exhibiting a ramping profile has been proposed as neurophysiological evidence for integration in DDMs (Gold & Shadlen, 2007). Finally, DDMs are easily extended to incorporate other reasonable features that can explain observations that do not match the basic DDM (Ditterich, 2006; Kiani, Hanks, et al., 2008; Ratcliff & Rouder, 1998; Ratcliff & Smith, 2004). However, recent studies have not only shown that perfect integration is not an optimal mechanism, especially under limited time (Durstewitz & Deco, 2008; Miller & Katz, 2010, 2013), but they have also undermined neurophysiological evidence for such integration (Katz, Yates, et al., 2016; Latimer, Yates, et al., 2015; Stanford, Shankar, et al., 2010; Thura, Cos, et al., 2014).

Traditionally, most of the experimental studies on SAT have explored this tradeoff by emphasizing the speed or accuracy—through instructing subjects to “be as fast as possible” versus to “be as accurate as possible”—or alternatively by fixing the total available time to make successive decisions (Bogacz, Wagenmakers, et al., 2010; Drugowitsch, DeAngelis, et al., 2015). Not only do such constraints rarely exist in real-world decision making, they are also likely to make subjects artificially favor one factor over the other. By contrast, in many real-life situations, potential reward outcomes and costs associated with different alternatives change over time, requiring the brain to dynamically integrate sensory and reward information to determine the proper time to terminate integration and commit to a choice. Nevertheless, the computational mechanisms underlying dynamic combination of sensory and reward information are largely unknown. It is also unclear whether this combination is affected by limitations in the integration of sensory information over time.

In order to address these questions, here we utilized a series of novel experiments in which we manipulated time pressure (i.e., the cost of time) over a wide range of values. Crucially, we implemented various levels of time pressure to closely probe sensory integration over time with different time intervals while subjects made decisions based on sensory information alone or by dynamically combining sensory and time-varying reward information.

## Results

To summarize our approach, we performed three experiments to study how sensory and reward information are dynamically combined over time. In Experiment 1, subjects made decisions about the dominant color in a patch of colored dots based on sequentially presented information within a limited time window (explicitly represented by a peripheral arc) to obtain reward points (Fig. 1A). In Experiment 2, subjects performed the same task while the peripheral arc represented the size of the potential gain (for a correct decision) or loss (for an incorrect decision), requiring subjects to dynamically combine sensory and reward information over time (Fig. 1B). In Experiment 3, a central bar representing the probability of reward (from 0.5 to 1.0) replaced the patch of colored dots (Fig. 1C). This central bar increased according to the SAT function for a given subject, while the peripheral arc decreased over time similarly to Experiment 2, enabling us to mimic Experiment 2 without a need to integrate sensory information over time in order to estimate the probability of reward. Therefore, Experiments 1 and 3 could serve as two controls for dynamic combination of sensory and reward information studied in Experiment 2 since Experiment 1 does not involve dynamic combination of sensory and reward information, and Experiment 3 does not require the integration of sensory information over time.

**Figure 1.**
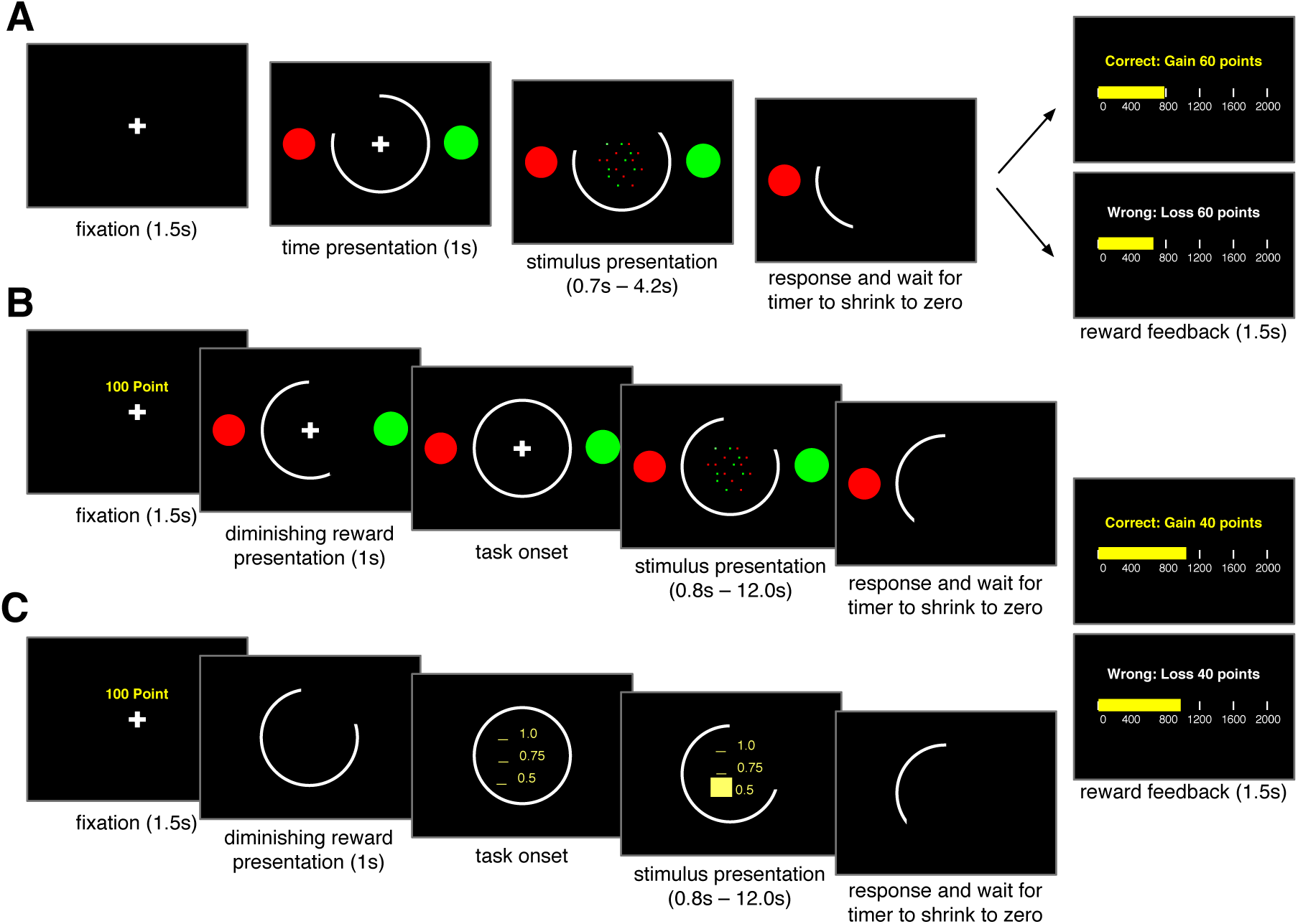
Schematic of experimental paradigms. (**A**) The timeline of the sequential-sampling task used in Experiment 1. After fixation, two color targets and a peripheral white arc appeared on the screen. The peripheral arc represented the available time. Subsequently, the fixation cross was replaced by green and red dots, which appeared randomly around the center of the screen while the size of the peripheral arc started to decrease. After deciding the dominant color of the dots and submitting their response, subjects had to wait until the timer reached zero, so there was no advantage in responding early. Finally, reward feedback was displayed, and the local accumulated reward points were updated accordingly. (**B**) The timeline of Experiment 2 in which the peripheral arc represented the available reward. Following the appearance of the fixation cross and information about the maximum reward (50 and 100 points), subjects were presented with a diminishing arc informing them about the rate of reward decrease on a given trial. The rest of the trial was similar to Experiment 1. (**C**) The timeline of the time-dependent gambling task used in Experiment 3. Similar to Experiment 2, after the fixation cross and maximum reward points, subjects were presented with a diminishing arc informing them about the rate of reward decrease on each trial. A central yellow bar represented the probability of reward (between 0.5 and 1.0), while the peripheral arc indicated the amount of reward. The subjects’ task was to accept the presented time-dependent gamble at any point during a trial, and reward feedback (reward or no reward with a magnitude depending on the time of response) was provided at the end of each trial.

Finally, in order to capture sensory integration under time pressure, we simulated choice behavior and reaction time (RT) during Experiment 1 using seven different computational models (six versions of the DDM and an attractor model with urgency signal; see Materials and Methods). In addition, we also compared the pattern of observed and optimal RTs during Experiments 2 and 3 to investigate combination of sensory and reward information under time pressure.

### Time pressure reduces performance during sensory integration

In Experiment 1, we implemented six different levels of time pressure by imposing hard deadlines (from 0.7s to 4.2s with 0.7s increment). Importantly, the level of time pressure was randomly assigned on each trial and signaled by the size of the peripheral arc. We first computed the average performance (i.e., the probability of making a correct response) and the mean RT for each level of time pressure. Overall, subjects’ performance gradually improved when they were given more time to respond but plateaued when the deadline exceeded 2.4s (Fig. 2A).

**Figure 2.**
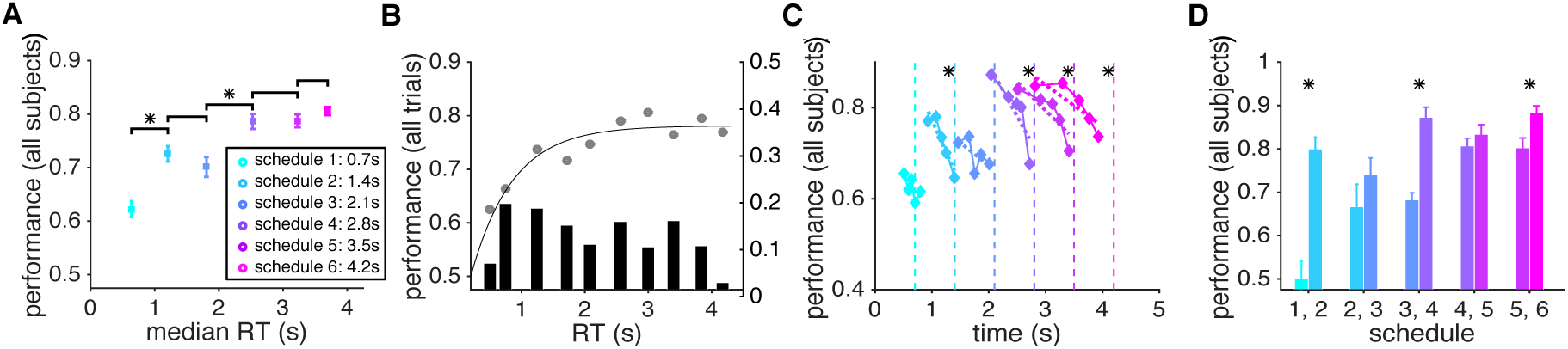
Performance of sensory integration over time is reduced by the leak in integration and with greater time pressure. (**A**) Average speed-accuracy tradeoff across subjects. The average performance (probability of a correct response) is plotted as a function of the average median response time (RT) across all subjects, separately for trials with a given level of time pressure. Schedule 1 to 6 correspond to trials with a deadline at 0.7s, 1.4s, 2.1s, 2.8s, 3.5s, and 4.2s, respectively. Averages for different schedules are similarly color coded in all panels. An asterisk above a line indicates that the difference between the performance values for the corresponding schedules was significant (sign test, *p <* 0.05). (**B**) Performance is plotted as a function of RT binned in ten equal intervals across all subjects. Performance increased with longer RT but plateaued at ∼2.25s, indicating the leak in integration of sensory information over time. Note that revealed dots stayed on the screen for 2.0s. The solid curve shows the fit using a modified Weibull function (Eq. 4 in Materials and Methods). The histogram shows the distribution of RT across the ten bins. (**C**) Performance reduced as the deadline approached. For each level of time pressure, we divided RTs into five bins and computed the probability of a correct response for trials in a given bin. The dashed lines indicate deadlines, and the asterisk next to each line indicates that performance for the corresponding schedule was significantly worse for longer RT (significant regression slope, *p <* 0.05). (**D**) Even with equal integration time, performance was worse under greater time pressure. Plotted is the average performance on RT-matched trials (based on mean) with adjacent levels of time pressure. The asterisk above each pair of histograms indicates that the difference between the performance values for the corresponding schedules was significant (sign test, *p <* 0.05).

During a schedule with the highest level of time pressure (schedule 1 with a time window equal to 0.7s), subjects were correct in approximately 62% of the trials. Subjects’ performance improved and plateaued at ∼78% with less time pressure and longer integration time (schedules 4, 5, and 6). Importantly, subjects used most of the available time in a given schedule for integration of sensory information—the median ratio of response time to the available time in all schedules was greater than 0.8s. Consistent with previous findings on the pattern of RT during different perceptual tasks (Luce, 1986; Ratcliff, 1978; Roitman & Shadlen, 2002), we found that compared to the incorrect trials, correct ones had shorter RTs under most levels of time pressure (Fig. 3A).

**Figure 3.**
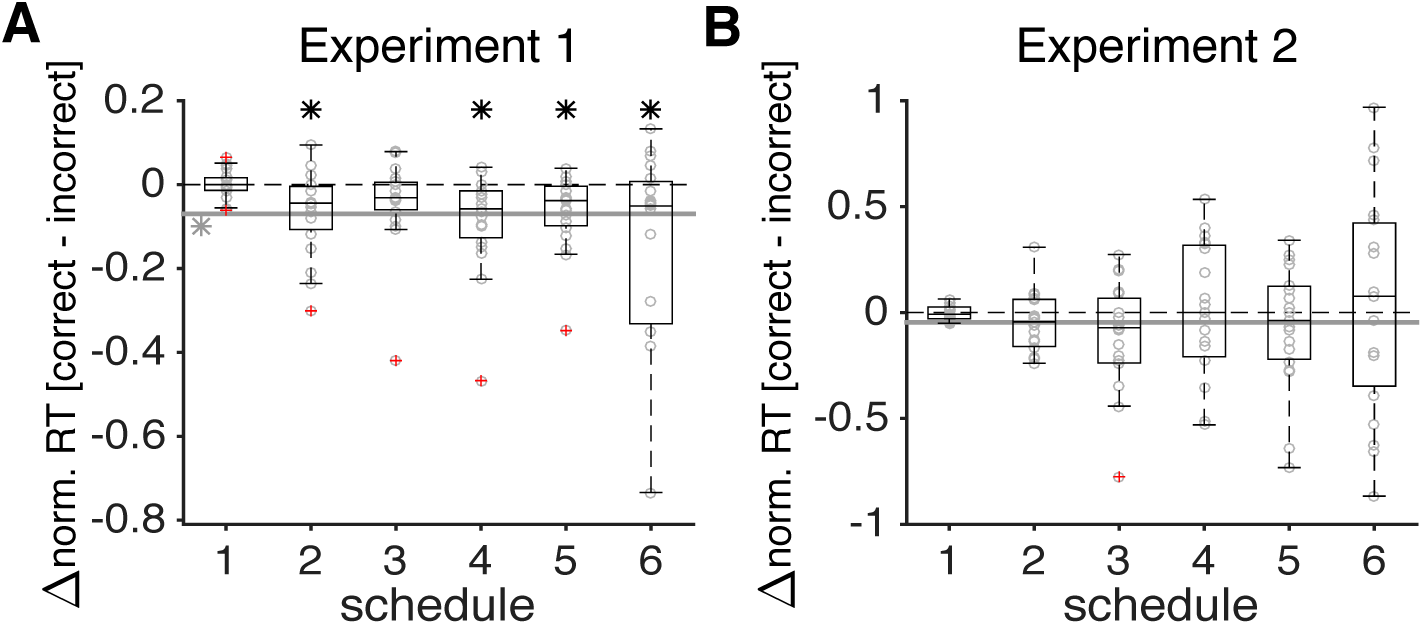
Response time was shorter on correct than incorrect trials in Experiment 1 but not in Experiment 2. (**A**) Plotted is the difference in normalized RT between correct and incorrect trials across all subjects in Experiment 1. The solid gray line shows the average difference over all schedules, and the gray asterisk next to it indicates that this difference was significantly different from 0 (sign test, *p* < 0.05). Each black asterisk shows that the difference in RT between correct and incorrect trials for a given schedule was significantly different from 0 (sign test, *p* < 0.05), and the outliers are indicated by circles with red crosses. (**B**) The same as in (A) but for Experiment 2. In Experiment 2, the difference in the normalized RT between correct and incorrect trials was not different from 0 in any schedule or in the average data.

In our experiments, the rate at which new samples (red or green dot) appeared on the screen (i.e., the sampling rate) was 20 Hz, and each sample stayed on the screen for two seconds or until the trial ended. The number of dots on the screen increased and reached 40 dots two seconds after the beginning of each trial and remained at this number throughout the rest of the trial. This design allowed us to study how sensory integration over time could be affected by a leak in integration since any imperfect integration of sensory information over time has a more significant effect on performance after this initial two-second period. Examining performance across different levels of time pressure, we found that performance reached its peak at around 2.25s and did not improve with longer RT even up to 4.0s (Fig. 2B). This lack of improvement indicates that instead of the entire sample history, only the most recent samples influenced choice in trials with long RT.

To directly test whether the most recent samples influenced choice more strongly, we used a linear logistic regression model to predict choice based on the sensory evidence (proportion of red dots) presented in the time intervals preceding a given choice. We found that the normalized weight of sensory evidence on choice to be the largest for dots appearing in time bins immediately preceding the choice and this weight decreased to zero on a timescale that was shorter than the deadline in a given schedule (Fig. 4A). Moreover, the decrease in the weight of evidence on choice was stronger for schedules with higher time pressure (schedules 1 to 4) indicating the direct influence of time pressure on sensory integration. Overall, these results illustrate that more distant samples have a weaker influence on subjects’ decisions, are more consistent with leaky integration of sensory information over time.

**Figure 4.**
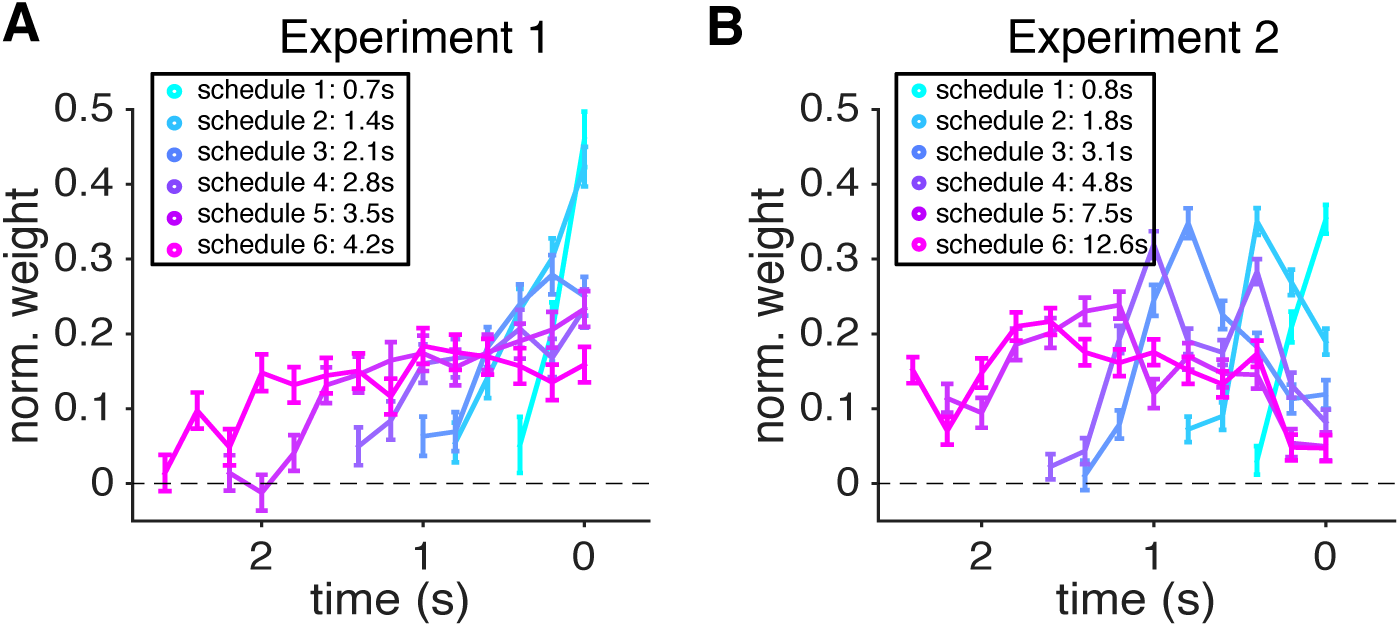
More recent samples had a stronger influence on choice in Experiment 1 but not in Experiment 2. The estimated weights of sample on choice are plotted as a function of time for Experiments 1 (A) and 2 (B), separately for different schedules. In Experiment 1, the influence of sensory evidence on choice dropped to zero on a timescale much shorter than the deadline in a given schedule. In Experiment 2, the most recent sensory evidence did not have the maximal influence on choice except in schedule 1.

Although the average psychometric function (Fig. 2A) indicates that longer RT led to better performance to some extent, reflecting the SAT, it does not reveal how approaching the deadline (i.e., time pressure) might affect performance. Perfect integration and a fixed threshold for making decisions predict that performance should improve when subjects respond closer to the deadline simply because having more time allows for accumulating more information, and subjects had no incentive to respond until just before the deadline. To test this prediction, we divided RTs into five equal bins and computed the probability of a correct response in each bin separately for each schedule. We found that in most schedules, the performance significantly dropped as subjects responded closer to the deadline (Fig. 2C). This happened despite the fact that subjects actually spent more time accumulating information before making a decision, indicating that the leak in sensory integration could be amplified by proximity to the deadline perhaps via an urgency signal (Cisek et al., 2009).

In order to test this possibility and reveal mechanisms underlying these observations, we fit the average performance and RT in Experiment 1 using various models and simulated behavior of these models using the estimated parameters (see Materials and Methods). These fitting and simulations illustrated that among all the DDMs we used, the DDM with urgency and leak (Fig. 5F) can most closely replicate the subjects’ performance shown in Figure 2, whereas the DDM with collapsing boundary (Fig. 5C) and the DDM with urgency (Fig. 5E) could capture the main features of our experimental data (Fig. 5). Our experimental findings are also consistent with a decision process based on stochastic transitions between discrete states modulated by an urgency signal (Fig. 5G). In this model, approaching the deadline causes more random transitions (due to an “urgency” signal; Miller & Katz, 2013) that can diminish performance closer to the deadline. Our simulations of the models also showed that although the basic DDM (Fig. 5A) and the DDM with leak (Fig. 5D) do not exhibit a decline in performance prior to the deadline, this phenomenon can be captured by the DDM with variant drift rates and the DDM with collapsing boundary (Fig. 5B-C and Supplementary Figure 1B-C). This happens in the DDM with variant drift rates because longer RTs more often occur on trials with smaller drift rates and due to a reduction of the threshold over time in the DDM with collapsing boundary. Together, these results illustrate that only the DDM with urgency and leak and the attractor with an urgency signal can most closely capture the experimental data shown in Figure 2.

**Figure 5.**
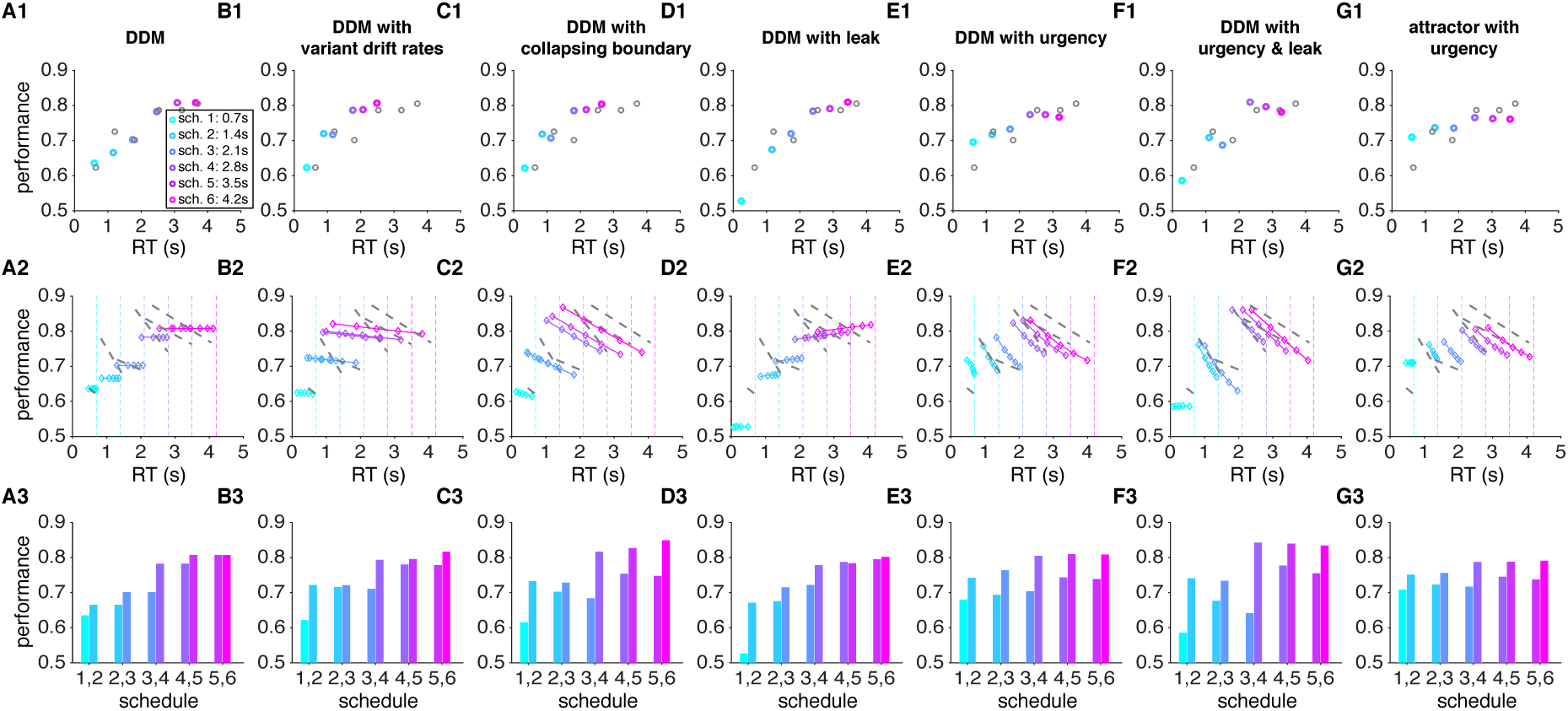
Simulations of experimental data in Experiment 1 using various models. Model simulations reveal that our data are more compatible with either leaky integration of sensory information with an additional urgency signal or sensory integration based on stochastic transitions modulated by an urgency signal. Results for different models are presented on different columns, while panels 1 to 3 of each column plot the average speed-accuracy tradeoff for a given level of time pressure (panel 1); performance as function of proximity to the deadline (panel 2); and the average performance on RT-matched trials (based on mean) with adjacent levels of time pressure (panel 3). The parameters for each model are determined based on the best fit of data in Figure 2C. For these simulations and fits, trials in which the decision boundary was not crossed before the deadline were removed from the analysis. The gray circles in the first column and dashed lines in the second column show the experimental results and linear regression fits of performance as a function of RT, respectively. Models include: basic DDM (A); DDM with variant drift rates (B); DDM with collapsing boundary (C); DDM with leak (D); DDM with urgency (E); DDM with urgency and leak (F); and attractor model with urgency (G). Conventions are similar to Figure 2.

To compute the effect of time pressure on sensory integration independently of the integration time, we next compared performance between adjacent levels of time pressure on trials with similar RT. By controlling for RT, we could directly measure the effect of time pressure on sensory integration. Accuracy was reduced under greater time pressure even when participants spent the same amount of time accumulating information and making a decision (Fig. 2D). Moreover, this effect was stronger for adjacent schedules with greater time pressure and gradually decreased as subjects were given more time to respond (less time pressure). Simulations of various DDMs revealed that lower accuracy for higher time pressure in RT-matched trials can be captured by assuming a lower threshold for schedules with higher time pressure even in the basic DDM. This effect, however, can be amplified with additional mechanisms such as an urgency signal or similarly a collapsing boundary (Fig. 5 and Supplementary Figure 1).

Overall, simulation results and data in Experiment 1 reveal a leak in sensory integration over time and are consistent with decision making based on stochastic transitions between states favoring alternative choices. Moreover, these results demonstrate that time pressure has a negative impact on performance irrespective of the integration time, and this effect becomes stronger as the response time approaches the deadline.

### Dynamic combination of sensory and reward information

In Experiment 2, subjects performed the same task as in Experiment 1 except that potential gain/loss for making a correct/incorrect decision continuously decreased with time at various rates (corresponding to different levels of time pressure). Overall, similar to what we observed in Experiment 1, performance improved with time (Fig. 6A) and under smaller time pressure (Fig. 6D). However, subjects used a smaller fraction of the available time in Experiment 2 and this effect was enhanced for schedules with lower time pressure (Fig. 6A). The probability of being correct was ∼60% in trials with the highest level of time pressure (schedule 1) and improved to ∼77% in schedules with smaller time pressure (Fig. 6A). In contrast to Experiment 1, however, the performance reached its maximum value with much less integration time (∼1.25s in Experiment 2 versus ∼2.25s in Experiment 1; Fig. 6B). That is, subjects reached the same level of performance as in Experiment 1 with less accumulated information (i.e., observing a fewer number of dots). Moreover, unlike Experiment 1, subjects’ performance did not drop as they responded closer to the deadline (except for schedule 3; Fig. 6C). Interestingly, when examining the impact of sample history on choice, we found that the normalized weight was not maximal right before choice and did not decrease immediately (Fig. 4B). This indicates that subjects stopped integrating new sensory evidence right before committing to a choice, perhaps due to enhanced processing of reward signals. Together, these results suggest that decision processes employed during the dynamic combination of sensory and reward information differs from those used during pure sensory integration.

**Figure 6.**
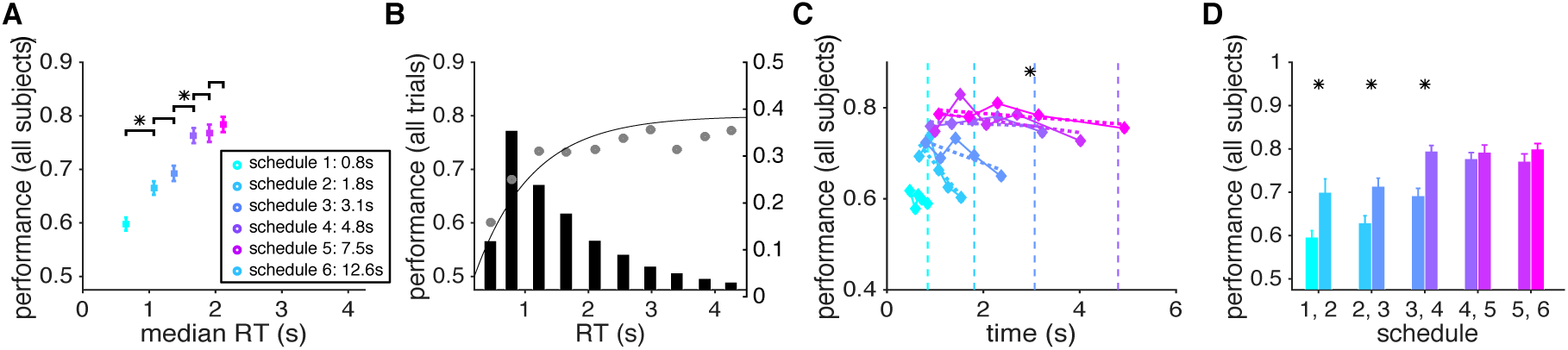
Dynamic combination of sensory and reward information was affected by time pressure and revealed that subjects were sensitive to imperfect sensory integration. (**A**) Subjects improved their performance with longer integration time, and their performance reached a plateau in less than 2.0s. The average performance (probability of a correct response) is plotted as a function of the average median response time (RT) across all subjects, separately for trials with a given level of time pressure (the average available time for each schedule is noted in the legend). Conventions are the same as in Figure 2. (**B**) Performance increased with longer RT but plateaued at ∼1.25s. Performance is plotted as a function of RT binned in ten equal intervals. The solid curve shows the fit using a modified Weibull function (Eq. 4 in Materials and Methods). The histogram shows the distribution of RT across the ten bins. (**C**) Performance did not improve and plateaued closer to the deadline (except for schedule 3 where it declined closer to the deadline) for a given level of time pressure. For display purposes, only four of the six deadlines are shown. (**D**) Even with equal accumulation time, performance was reduced with more time pressure. Plotted is the average performance on RT-matched trials (based on mean) with adjacent levels of time pressure.

We also tested the effect of time pressure on dynamic sensory-reward integration independently of the integration time by comparing performance between adjacent levels of time pressure on trials with similar RT. We found that similar to Experiment 1, performance was worse under greater time pressure even when subjects spent the same amount of time accumulating information and making a decision. However, the differences in performance between adjacent levels of time pressure were smaller than those observed in Experiment 1 and were only present for high-pressure schedules (Fig. 6D). Overall, time pressure implemented via decreasing reward value had less of a negative impact on performance than time pressure via a hard deadline as in Experiment 1.

To summarize, these results suggest that decision processes involved in dynamic combination of sensory and reward information are different from those required for integration of sensory information over time. More specifically, during dynamic combination of sensory and reward information, the brain seems to constantly monitor/evaluate sensory integration (or stochastic transitions in the state of the decision process) in order to determine when to terminate this process before the leak in this integration hinders improvement in performance. In other words, it is possible that by perceiving a leak in sensory integration, subjects were able to more quickly terminate this process and therefore reduce the cost of time.

### Response time depends on perceiving improvement in performance

In Experiment 2, we created time pressure by decreasing the size of potential reward over time. To maximize rewards, subjects had to appropriately balance the cost (the decreasing reward) and benefit (increasing probability of making a correct response) of accumulating information over time in order to determine the right time to make a choice.

To predict the optimal RT, we used a simple model in which an ideal decision maker integrates their speed-accuracy tradeoff (SAT) and the decreasing reward function to compute expected gain over time (Fig. 7A). The optimal decision strategy is to stop integration and respond at the time that yields the maximum expected gain (Fig. 7B). This simple model is parameter-free and only uses individual subjects’ SAT function (from Experiment 1) to predict the optimal response time for each schedule. Critically, the model assumes that individual subjects have access to their SAT function from which they can compute expected gain over time in order to find the time point with maximum expected gain (EG). We used this model to predict the optimal RT for each subject in each schedule.

**Figure 7.**
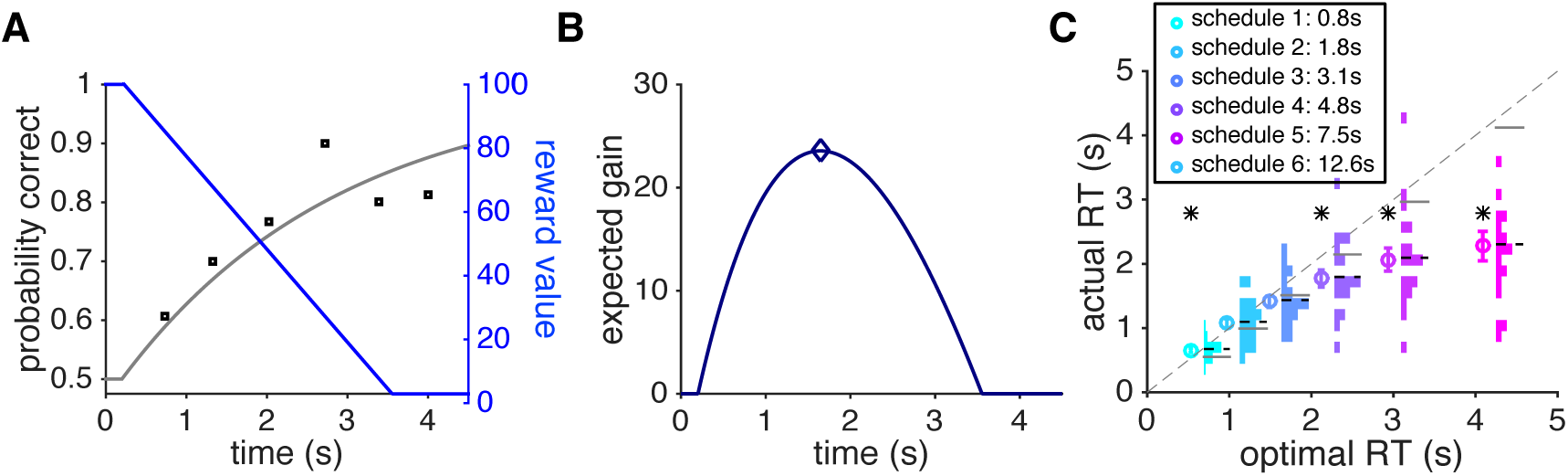
An optimal model of RT for dynamic sensory-reward combination under time pressure and comparison between the optimal and actual RT in Experiment 2. (**A**) Tradeoff between improved performance and less potential reward over time. The probability of a correct response (left axis) increases with longer integration time (the SAT curve in gray), while the reward harvested upon a correct response decreases over time (blue, right axis). Black circles show actual performance on trials with different levels of time pressure (based on data from an individual subject in Experiment 1), and the black line shows the fit based on the modified Weibull function (Eq. 1 in Materials and Methods). (**B**) The expected gain (EG), equal to the product of the SAT and reward function, is plotted as a function of time. An ideal decision maker uses the EG over time to pick the optimal RT. (**C**) Comparison between the actual RT and the optimal RT. Subjects systematically used less time than what is optimally prescribed to make decisions under time pressure for intermediate and slow schedules (schedules 3–6). Each circle marker represents the mean RT (± s.e.m.) of all subjects in a particular schedule against the optimal RT predicted by the ideal decision maker. Each histogram indicates the count distribution of median RT across subjects, and the horizontal dashed and solid lines represent the mean of each distribution and the optimal RT, respectively. The asterisk indicates statistical significance between the actual and optimal RT (sign test, *p <* 0.05).

Comparing the actual and optimal RT, we found that overall subjects spent less time on the integration of sensory information than prescribed optimally (Fig. 7C). Only when reward decreased most sharply (schedules 1 and 2) did subjects spend slightly more time than what is optimal to make a decision, perhaps due to failing to respond fast enough while processing different pieces of information. For other schedules, subjects spent less time than they should optimally, but this effect was only statistically significant for slower schedules (schedules 4 to 6; sign test, *p* < 0.05).

In Experiment 2, subjects faced a tradeoff between improving accuracy and diminishing reward on a correct response. This means that subjects could stop integration if they did not perceive any improvement in performance over time since more time spent on integration only corresponds to losing more reward. Therefore, perceiving the lack of improvement in performance could explain why subjects stopped integration early and responded more quickly than prescribed optimally.

To test whether or not perceiving improvement in performance could partially account for the sub-optimality observed in Experiment 2, we ran a control task equivalent to the one in Experiment 2 but did not require the integration of information over time. In this control task (Experiment 3), subjects decided when to accept a time-dependent gamble with a decreasing reward magnitude and an increasing reward probability (time-dependent gambling task; Fig. 1C). Importantly, the increase in the probability of reward over time was based on the subject’s performance in Experiment 1 (the SAT function), whereas the decrease in the magnitude of reward was identical to schedules in Experiment 2. Therefore, assuming that a subject’s SAT remains the same in both tasks and that the subject has access to this function, this task fully mimics the dynamic sensory-reward integration under time pressure in Experiment 2 without the need to integrate sensory information over time. Moreover, in the control task, the change in performance was unaffected by time pressure due to its proximity to the deadline.

We found that during gambling under time pressure, response time to accept a gamble showed a different pattern of deviations from the optimal RT observed in Experiment 2. For all but the lowest level of time pressure (schedule 6), subjects were slower than or close to the optimal RT (Fig. 8A). Moreover, subjects were overall slower in responding in Experiment 3 than in Experiment 2 (Fig. 8B) except for schedule 1 (the fastest schedule). In Experiment 3, the probability of reward increased over time based on each subject’s SAT function estimated in Experiment 1. In contrast, in Experiment 2, competition between processing the sensory information (red and green dots) and the cost of time (remaining reward) could mitigate sensory integration and thus, hinder the subjects from perceiving an improvement in performance over time. Together, these results suggest that faster-than-optimal RT during dynamic sensory-reward combination (Experiment 2) was partially caused by subjects’ perception of no improvement in their performance over time, which we ascribed to the leak in sensory integration enhanced by time pressure.

**Figure 8.**
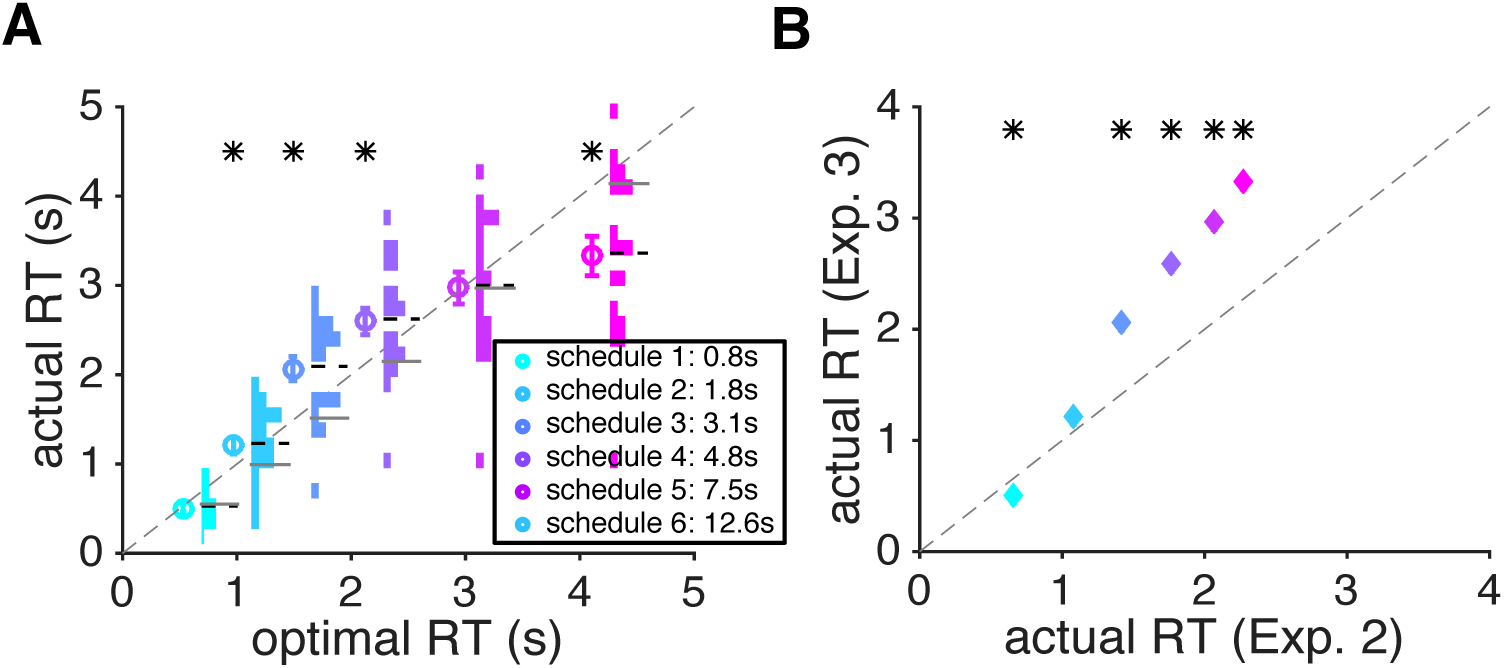
Different patterns of response time between the time-dependent gambling task (Experiment 3) and dynamic combination of sensory and reward information (Experiment 2). (**A**) Comparison between the actual RT and the predicted optional RT in Experiment 3. Conventions are the same as in Figure 7C. (**B**) Comparison between the average RT in Experiments 2 and 3. Overall, subjects showed larger RT in Experiment 3 for slower schedules. The asterisk indicates statistical significance between actual and optimal RT in panel A, and actual RT between Experiments 2 and 3 in panel B (sign test, *p<* 0.05).

In addition to the leak in sensory integration, loss aversion could also contribute to early decision-making termination and the observed suboptimal response time. At each point during a trial, subjects could compare the marginal gain in the probability of being correct with the marginal loss in the reward and then could terminate accumulation if the marginal loss exceeded gain (otherwise to continue integration). For a given level of time pressure, the marginal loss is constant across time, whereas the marginal gain decreases with time since the SAT function is concave. By magnifying the marginal loss, loss aversion would force subjects to respond earlier resulting in faster-than-optimal response times across all schedules. However, this mechanism would predict a similar response time pattern for Experiment 3, which is not what we found.

### Larger initial reward value improves dynamic sensory-reward combination

In Experiments 2 and 3, we used two different initial reward values for each level of time pressure. The optimal model predicts that for a given level of time pressure the optimal RT is independent of the initial reward value. In contrast, dynamic combination of sensory and reward information predicts that actual improvement in performance and the amount of decrease in reward should determine RT.

To test these contrasting predictions, we computed RT separately for the two initial reward values and each schedule. Contrary to the prediction of the optimal model, the initial reward value had a small but significant impact on subjects’ RT depending on the level of time pressure. Under greater levels of time pressure (schedules 1, 2, and 3), subjects spent more time when the initial reward value was smaller (Fig. 9A). In contrast, when facing lower levels of time pressure (schedules 4, 5, and 6), a smaller initial reward value produced shorter RT. Furthermore, we also found that modulations of RT by the initial reward value made the behavior closer to optimal when the initial reward value was larger (Fig. 9B), with the RT being significantly closer to optimal in four out of six schedules. As aforementioned, the diminishing reward arc was presented on the screen before the task began in each trial (Fig. 1B–C), and thus, the uncertainty associated with the perception of reward size and how it changed over time was similar between the two conditions.

**Figure 9.**
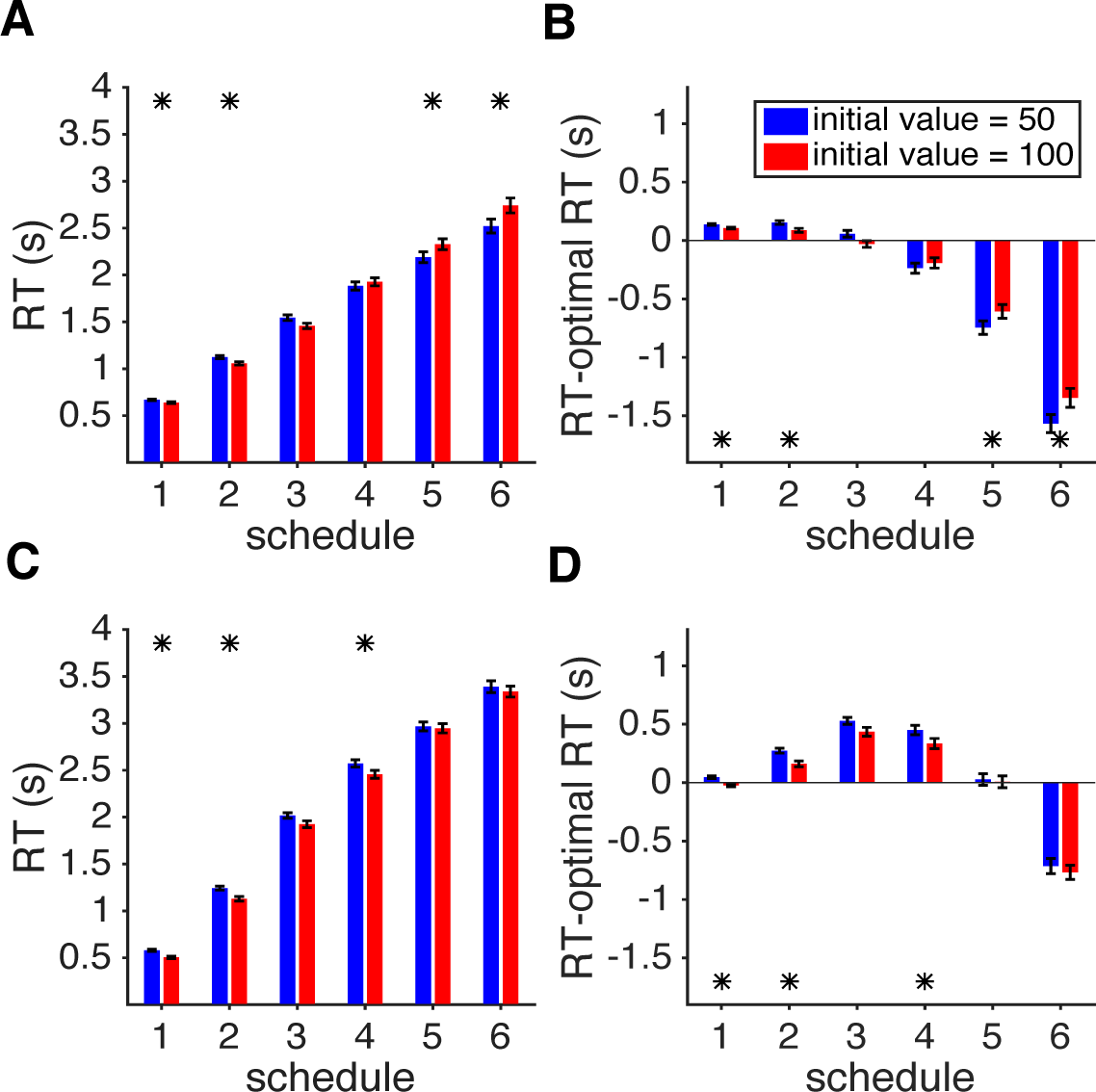
The effect of initial reward value on dynamic combination of sensory and reward information. (**A**) For schedules with greater levels of time pressure, a larger initial reward value (red) resulted in faster RT (schedules 1 and 2) than a smaller initial reward value (blue), but the opposite was true for schedules with smaller levels of time pressure (schedules 5 and 6). Plotted is the median RT separately for each level of time pressure and initial reward value in Experiment 2. Error bars show the s.e.m. (**B**) The median of the difference between the actual RT and optimal RT is plotted separately for each level of time pressure and initial reward value in Experiment 2. Overall, a larger initial reward value made RT closer to optimal. The asterisk indicates a significant difference in RT between the two initial reward values (sign rank test, *p* < 0.01). (**C, D**) The same as in (**A, B**) but for Experiment 3. Overall, RT was larger for trials with a smaller initial reward value in all schedules, indicating that subjects waited longer to reach a higher probability for accepting an initially smaller reward.

In contrast to Experiment 2, subjects responded slower on trials with a smaller initial reward value in all schedules of Experiment 3, but the difference in RT was significant for only three schedules (Fig. 9C). This indicates that when sensory integration was not required (and thus there was no leak), subjects waited longer on average to improve the probability of obtaining an initially smaller reward. Moreover, we did not find any evidence that the actual RT was closer to the optimal RT when the initial reward value was larger in Experiment 3 (Fig. 9D). Comparison of results between Experiments 2 and 3 suggests that a larger initial reward value can only improve dynamic combination of sensory and reward information by slowing down RT and making it closer to the optimal RT.

### The effects of previous outcome and reward schedule on response time

To further study the potential differences in decision processes between the three experiments, we next examined the trial-by-trial adjustment of response time. More specifically, we tested two factors that could contribute to the amount of time spent on integration: reward outcome and the level of time pressure on the preceding trial.

First, we examined whether RT on a given trial was affected by reward outcome (correct/incorrect or gain/loss received after a choice) on the preceding trial. In Experiment 1, subjects became slower (larger normalized RT) following correct compared to incorrect trials, but this effect was significant for only two schedules (schedules 4 and 5; sign rank test, *p* < 0.05; Fig. 10A). In contrast, there was no evidence for modulation of RT by reward outcome on the preceding trials in all schedules of Experiment 2 (Fig. 10B). Moreover, there was a large difference between RT following rewarded and unrewarded trials in Experiment 3, such that subjects waited longer to increase the probability of obtaining a smaller reward following rewarded trials (Fig. 10C). Therefore, in both Experiments 1 and 3, subjects tended to respond slower on trials following rewarded trials (perhaps to increase their chances of getting another reward), but this was not observed in Experiment 2.

**Figure 10.**
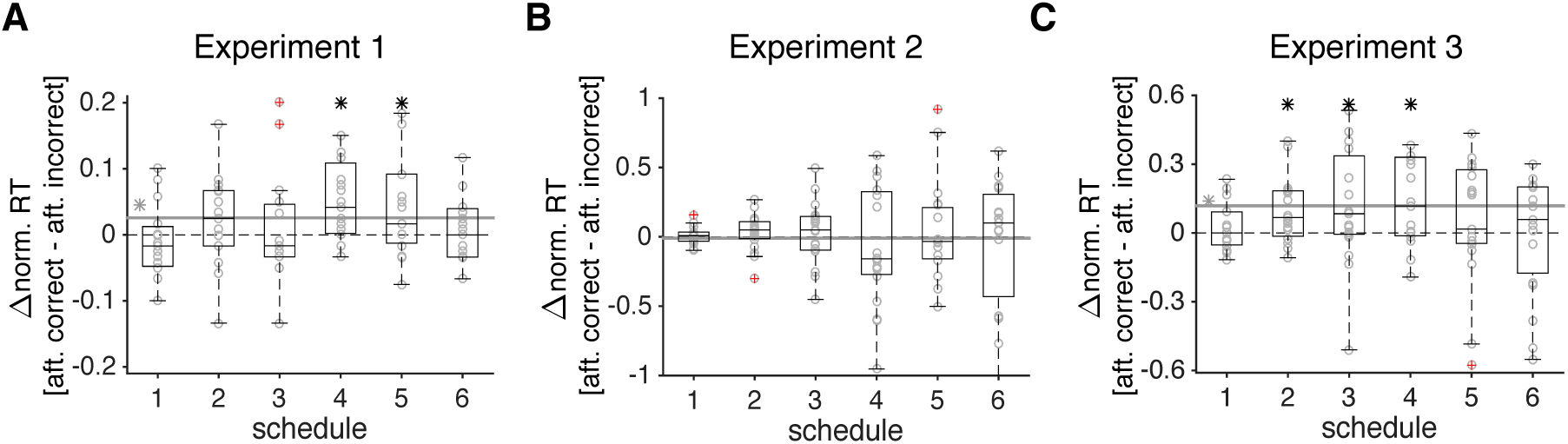
The effects of reward outcome on RT in the subsequent trial. (**A**) The bar plot shows the difference in median RT after correct and incorrect trials in Experiment 1, and the gray circles show the difference in median RT for individual subjects. The solid gray line shows the average difference over all schedules, and a gray asterisk next to it indicates that this average was significantly different from 0 (sign test, *p* < 0.05). A black asterisk indicates that the difference in RT for a given schedule was significantly different from 0 (sign test, *p* < 0.05), and outliers are indicated by circles with a red cross. (**B**, **C**) The same as in (A) but for Experiments 2 and 3, respectively.

Together, these results illustrate that RT was not affected by reward outcome on the preceding trials only in Experiment 2, which required dynamic combination of sensory and reward information. As mentioned above, this combination is modulated by the perception of leak in sensory integration on each trial (i.e. lack of improvement in performance), which can override any effect of preceding reward outcome. In addition, unlike Experiment 1, subjects’ RT on correct trials did not differ from incorrect trials in Experiment 2 (Fig. 3B), indicating that the underlying mechanisms are different between pure sensory integration and sensory-reward combination.

Second, we tested whether the level of time pressure on a given trial had any effect on RT in the following trial in Experiment 1. We did not consider this analysis for Experiments 2 and 3 since in these two experiments, subjects knew the level of time pressure at the beginning of the trial and experienced the new level of time pressure before entering the task phase (Fig. 1B–C). We found that on schedules with moderate levels of time pressure (schedules 2 and 3), subjects responded slower following schedules with lower time pressure than greater time pressure (sign rank test, *p* < 0.05). This result illustrates that the previous level of time pressure may have a lasting effect on behavior—perhaps by adjusting the threshold—and indicates that the brain can adjust to time pressure with great fidelity.

## Discussion

Investigating how the brain integrates information over time has been a central focus of decision-making studies at both experimental and theoretical levels (Busemeyer, Wang, et al., 2006; Gold & Shadlen, 2001, 2007; Hernández, Zainos, et al., 2002; Ratcliff, 1978; Usher & McClelland, 2001). Time constraints and pressure often reduce the quality of decisions (Kocher & Sutter, 2006; Payne, Bettman, et al., 1993), whereas collecting a greater amount of information improves performance (Griffin & Tversky, 1992). Therefore, exploring the SAT can reveal temporal aspects of the decision-making processes. Although many studies have investigated the effect of time pressure on integration of sensory information (see Bogacz, Wagenmakers, et al., 2010), no study has considered time pressure over a wide range of timescales.

Traditionally, time pressure has been created by setting a fixed time constraint, for example by imposing a deadline that is one standard deviation below the mean response time on the task of interest (Benson & Beach, 1998; Maule, Hockey, et al., 2000; Ordonez & Benson, 1997). In contrast, we created time pressure in two different ways: 1) by limiting integration time by imposing a hard deadline using a wide range of intervals; and 2) by making reward a decreasing function of time using a wide range of decreasing rates. By having schedules with various levels of time pressure, we were able to examine sensory integration over a wider range of time intervals. By implementing a cost function—potential reward decreasing in value over time at different rates—we explored the dynamic combination of sensory and reward information under time pressure. To our knowledge, no study has investigated these issues before although previous studies have examined the effects of time pressure on performance during decision making (Benson & Beach, 1998; Busemeyer & Townsend, 1993; Kocher & Sutter, 2006; Maule, Hockey, et al., 2000; Ordonez & Benson, 1997; Payne, Bettman, et al., 1993) and characterized the dynamics of decision time when there was a tradeoff between time and performance (Battaglia & Schrater, 2007; Dean, Wu, et al., 2007; Juni, Gureckis, et al., 2016; Rieskamp & Hoffrage, 2008).

We found that as the deadline approached, subjects’ performance decreased even though they actually spent more time accumulating sensory information. Moreover, performance worsened under greater time pressure even when subjects used the same amount of time for accumulating sensory information. Simulations of various models revealed that these could occur due to an urgency signal pushing the subject to make decisions as the deadline approaches (similarly by decreasing the decision boundary over time) or smaller drift rates on trials with longer RTs. Divided attention between processing of the sensory stimulus and the remaining time, which could become more biased towards the latter as the deadline approaches, could also contribute to the observed decrease in performance with longer RT.

Although a lack of improvement in performance has been reported, the leak in sensory integration has not been clearly demonstrated in previous behavioral studies mainly due to limitations in the experimental designs (but see Carland et al., 2016). For example, Usher and McClelland (2001) reported performance saturation that they attributed to leaky integration. However, it has been argued that saturation of performance could happen during fixed-duration discrimination tasks if the subjects terminate integration before an imposed deadline since they are not allowed to choose when to respond (Kiani et al., 2008). In other words, lack of improvement may not be caused by leaky integration but rather by the timing in the termination of integration. To address this issue, the ideal design would present information sequentially and to let subjects freely choose when to make a response. Our experimental paradigm had both of these design features. Using this design, we found the effect of presented evidence on choice gradually decreases for more remote evidence indicating a leak in sensory integration over time.

In an attempt to shed light on the underlying mechanisms, we simulated the behavior of six different models –– six versions of DDM and an attractor model with urgency — and found that perfect integration cannot account for our experimental findings. Our simulations based on the fit of experimental data with each model illustrate that the best models for capturing the performance and RT data in Experiment 1 are the DDM with urgency and leak, and the attractor model with urgency. The DDM with trial-to-trial variability of signal intensity (the DDM with variant drift rates) can account for correct decisions to be faster than incorrect ones and for a decline in performance as the deadline approaches. However, this model cannot fully capture all aspects of our experimental data. We also note that the inter-trial interval was padded with the remaining time and thus, subjects had no incentive to respond before the deadline if they were not confident with their choice. In other words, on trials in which the signal intensity was weak, subjects could simply spend more time making a decision. Together, our results point to a leak in the integration of sensory information over time, which is further amplified by time pressure due to an urgency signal.

Our results are also compatible with decisions based on stochastic transitions between discrete attractor states, corresponding to lack of integration over time (Durstewitz & Deco, 2008; Katz, Yates, et al., 2016; Latimer, Yates, et al., 2015; Miller & Katz, 2010). For very fast schedules in Experiment 1, the decrease in performance before the deadline is compatible with transitions to the state favoring the opposite direction of sensory signal due to approaching the deadline. Interestingly, Miller and Katz (Miller & Katz, 2010; 2013) had shown that when internal noise is significant, decision making based on stochastic transitions can improve performance under limited time. In our experiment, schedules with long response times could increase the contributions of internal noise (e.g., due to alternation between different sources of inputs) such that decision making based on transitions between attractor states becomes more advantageous than integration (Durstewitz & Deco, 2008; Miller & Katz, 2010; 2013).

A few recent studies have investigated whether humans can select decision time by trading off speed and accuracy so as to maximize reward rate (Bogacz, Wagenmakers, et al., 2010; Drugowitsch, DeAngelis, et al., 2015). These studies focused on the SAT in the context of maximizing reward rate. They, however, did not consider the tradeoff between speed and accuracy that could take place outside the possibility of reward-rate maximization, which happens in many real-life decisions that are one-shot in nature. For example, a baseball player—when deciding whether and when to swing at a ball—most likely is not concerned about maximizing reward rate because how long it takes her to make such a decision does not affect the amount of time she has the next time she steps onto the batting field. Yet, in this particular batting opportunity, the tradeoff between speed and accuracy is evidently important to making the batting decision. This is the spirit and the kind of SAT problem our experiment tried to capture. In the context of our experiment, the decision problem in each trial is how much time to spend on integrating information over time given the cost function (decreasing reward over time). Finally, the overall length of the experiment and the number of trials were fixed in our experiments. Therefore, for the subjects, there was no incentive to respond faster or to derive a decision strategy in an attempt to maximize reward rate.

During dynamic combination of sensory and reward information, the amount of time subjects spent on sensory integration was overall less than the optimally prescribed RT, and the extent of deviation from the optimal RT increased as time pressure was reduced. Comparing the patterns of response times in Experiments 2 and 3 revealed that the suboptimal RT could be related to the perception of leak in sensory integration (i.e. lack of improvement in performance) enhanced by time pressure (absent in Experiment 3). These observations indicate that subjects not only perceived this leak but also incorporated it into their decision-making processes.

Perception of leak in sensory integration could involve metacognition (Deroy, Spence, et al., 2016), a second-order judgment that allows the decision maker to monitor the first-order response, such as deciding the dominant color in our Experiments 1 and 2. Previous research on metacognition suggests the involvement of the rostrolateral prefrontal cortex, the striatum, and the thalamus in metacognitive judgment (Baird, Smallwood, et al., 2013; Fleming, Dolan, et al., 2012; Fleming, Huijgen, et al., 2012; Fleming, Weil, et al., 2010; Hebart, Schriever, et al., 2014; Wu, Delgado, et al., 2015). Part or all of these networks could be involved in detecting and representing the perception of leak in sensory integration and possibly communicating this information to the decision circuit. Future experiments can test for a possible role of metacognition in dynamic sensory-reward combination.

Here, we did not explicitly model the integration of sensory and reward information over time using the DDMs because there are many ways in which sensory and time-dependent reward information could be combined in the framework of DDMs, and we did not have enough experimental data to test these models against each other. For example, a diminishing reward can be implemented by a decision boundary that decreases to zero at different rates (for different reward schedules) or with different drift rates. However, these two models can be hardly distinguished from each other, making it nearly impossible to determine the correct model based on our human data with limited number of trials. Due to this limitation, we only focused on the simulation of performance and response time in Experiment 1 and used our normative model to address the pattern of response time in Experiments 2 and 3. In comparison, two recent models for combination of sensory and reward information are either concerned with decision making in the presence of a fixed reward bias (Gao, Tortell, & McClelland, 2011) or consider reward values to depend on time because of the state of the subject during the task (Christopoulos & Schrater, 2015). Therefore, neither of these models can be used to simulate performance and reaction time in Experiment 2 of this study. Nevertheless, comparison of the predictions of our normative model for the pattern of RT between Experiments 2 and 3 revealed that the need for integrating sensory information over time resulted in shorter RT in Experiment 2.

We note that although mathematically equivalent, Experiments 2 and 3 can be different cognitively. More specifically, in Experiment 2, subjects had to make a perceptual judgment in order to receive a reward, whereas Experiment 3 did not involve any such judgment. Instead, information about probability of reward was explicitly revealed to the subjects in Experiment 3, and the task was to decide when to press a button to realize the time-dependent lottery. Compared with Experiment 2, Experiment 3 resembled more of a typical value-based choice task. It is therefore possible that some of the differences in the pattern of response time between Experiments 2 and 3 are due to differences between perceptual and value-based decision making. Moreover, Experiment 2 involved uncertainty due to noisy sensory information whereas in Experiment 3 the probability of reward was represented by the height of a bar and thus, the latter experiment involved risk only. Previous studies showed that humans are more risk averse when probability of reward is moderate to high (>35∼40%) in tasks where information about probability is explicitly presented (Kahneman & Tversky, 1979). Our findings in Experiment 3 are consistent with this finding: the fact that subjects generally exhibited slower RT in Experiment 3 is consistent with the notion that they were more risk averse in Experiment 3 than Experiment 2.

Finally, comparison of results between Experiments 1 and 2 revealed important differences between decision processes underlying pure sensory integration (Experiment 1) and dynamic sensory-reward combination (Experiment 2). First, during dynamic sensory-reward combination the subjects reached the same performance level of sensory integration under time pressure but with a shorter integration time. A similar phenomenon was observed in a beauty-contest game wherein time-dependent payoffs under high time pressure lead to significantly quicker decision making without reducing the quality of decisions (Kocher & Sutter, 2006). Second, unlike pure sensory integration, performance did not drop as the deadline approached during sensory-reward combination. Finally, RT neither differed between correct and incorrect trials nor depended on reward outcome in the preceding trial. These differences could have important implications for the neural mechanisms involved in dynamic sensory-reward combination.

More specifically, there are currently two general alternative mechanisms for combination of sensory and reward information (opportunity cost). First, these two pieces of information could be combined in one decision circuit that determines both the time to respond and a response. Based on previous studies on perceptual decision making, the posterior parietal and lateral prefrontal cortices could provide plausible circuits for performing dynamic combination of reward and sensory information (Gold & Shadlen, 2007; Heekeren, Marrett, et al., 2008). However, recent evidence has shown that pharmacological inactivation of lateral intraparietal cortex (area LIP) does not affect performance during perceptual decision making (Katz et al., 2016), undermining the causal role of this area in choice and calling for future investigation (Yates, Park, Katz, Pillow, & Huk, 2017). Other candidate areas are the ventromedial prefrontal and orbitofrontal cortices, which have been shown to represent the decision variable accumulated over time in value-based choice (Gluth, Rieskamp, et al., 2012). Alternatively, two separate circuits could be involved in signaling the cost of time and integration of sensory information over time. Observed differences in the relationship between RT and performance across our experiments are more compatible with this possibility. However, future experiments are required to distinguish between the two alternatives.

## Materials and Methods

### Ethics statement

The study was approved by the Institutional Review Board at the Taipei Veterans General Hospital. Before the experiment, all subjects gave written consent to participate in the study.

### Subjects

Nineteen subjects participated in the study (nine males, mean age = 23.2 years). No participants had a history of psychiatric or neurological disorders.

### Overview of the experiments

To study how humans dynamically combine sensory and reward information and to implement various levels of time pressure on sensory integration, we designed a novel perceptual decision-making task in which subjects had to decide the dominant color of a sequentially presented patch of dots. Each subject performed a pre-test task and three experiments—the main interest of the study was the three experiments performed and not the pre-test task (see *pre-test task* below for more details).

For the pre-test task and the first two experiments (Fig. 1A–B), subjects were instructed to determine the dominant color of an imaginary box containing colored dots (green and red, 100 dots in total). In each trial, the subjects were first presented with information about how much time they had (pre-test and Experiment 1, Fig. 1A)—indicated by the size of the peripheral arc—or the magnitude of reward value (Fig. 1B–C), which was also indicated by the size of the peripheral arc. In Experiments 2 and 3, the arc subsequently decreased in size for 1.0s in order to inform the subject about the rate at which the reward would decrease over time in that trial (the rate could change from trial to trial). This phase was followed by a brief fixation period before entering the task phase of the trial. Following the fixation period, colored dots were drawn randomly from the box (with replacement) and were sequentially revealed. Every 50.0ms a new dot was shown (sampling rate equal to 20 Hz), and it stayed on the screen for 2.0s. The dots were presented within an aperture invisible to the subjects (5.25° visual angle). The luminance of the red and green colors was matched.

Subjects were instructed to indicate the dominant color of the dots by choosing the corresponding color target (red or green discs that were presented on the periphery) using the left or right arrow key. They could indicate a decision at any point in time after the onset of the sample (dots) presentation. Once a color was selected, both the dots and the chosen target disappeared. However, the arc continued to decrease in size and reached zero before the presentation of reward feedback. That is, the subjects had to wait for the arc to decrease to zero regardless of when she or he made a response. Therefore, there was no incentive (in terms of the time) for the subjects to respond faster, since the total length of the experiment and the number of trials were independent of the subjects’ performance. The local accumulated reward points (shown by a golden bar at the bottom of the screen; Fig. 1) were updated after each reward feedback and were reset when they reached 2000 points. This reset was accompanied by the message, “Congratulations!! $40 just deposited to your account,” which appeared on the screen and indicated to the subject that they had secured a specific amount of money (2000 points equaled 40 National Taiwan Dollars, or NTD; 1 US dollar equals 32 NTD).

### Manipulation of time pressure

In Experiment 1, time pressure was created by imposing a hard deadline for each trial. Subjects had to indicate their decision about the dominant color within a fixed time window (indicated by the length of a peripheral arc) in order to obtain a fixed reward (for correct responses) or endure a fixed loss. The time windows were selected from six possible values of 0.7s, 1.4s, 2.1s, 2.8s, 3.5s, and 4.2s, corresponding to schedules 1 to 6, respectively. The time window (i.e., schedule) was pseudo-randomly assigned in each trial with the constraint that all of the schedules were equally presented. In Experiments 2 and 3, different levels of time pressure were created by making the reward feedback decrease with time at different rates. The amount of gain or loss subjects received after a response was represented by the size of the white peripheral arc that decreased over time at different rates corresponding to different levels of time pressure. The reward schedule was pseudo-randomly assigned on each trial. The collected points were converted to actual money at the end of each experiment. Similar to Experiment 1, subjects were free to respond at any given point during a trial.

### Pre-test task

Each subject performed a pre-test task. The goals of this were: 1) to familiarize subjects with the decision task in Experiment 1; and 2) to estimate the correct parameter for the task in Experiments 1 and 2. The main parameter to estimate was a suitable dot ratio (i.e., the ratio of the dominating dots to the dominated dots) that allows the measurement of the speed-accuracy tradeoff (SAT) under various levels of time pressure. Based on a previous pilot study, we used 3 possible dot ratios: 60:40, 57:43, and 54:46. To measure the SAT for each dot ratio, we used 3 different time-window values: 0.7s, 2.1s, and 3.5s. Therefore, we had a 3 (dot ratio) × 3 (time window) factorial design consisting of 9 blocks (each with 36 trials) and 324 trials in total. For each block, there was a unique dot ratio. The order of dot ratios across the 9 blocks was always fixed—starting at 60:40, followed by 57:43 and 54:46 before going back to 60:40 again to repeat the sequence. Within a block, the time windows were randomly assigned in each trial. In each trial, subjects could submit their response about the dominant color at any point within the time window and would gain 60 points for a correct response or lose 60 points for an incorrect response (Fig. 1A). If the subject did not respond before the time expired, they would lose 60 points. Based on the pre-test, we picked the dot ratio for each subject such that under this ratio the subject exhibited reasonable SAT. For most subjects, this ratio was 57:43. This ratio then became the single fixed dot ratio used in Experiments 1 and 2.

### Experiment 1: Integration of sensory information under time pressure

In this Experiment, subjects performed the same task as in the pre-test session except six different time windows (0.7s, 1.4s, 2.1s, 2.8s, 3.5s, and 4.2s) were used with a fixed dot ratio. There were ten blocks, each with 36 trials, of 360 total trials. In each block, subjects faced all possible time windows equally, so each time window was presented six times.

### Estimating the speed-accuracy tradeoff function

For each subject, we used the data from Experiment 1 to estimate the SAT function, which was then used to tailor parameters for Experiments 2 and 3. Recall that there were six different time windows in Experiment 1, each of which was presented in 60 trials. For each time window, we calculated the mean response time (RT) and the proportion of correct trials. These data were then used to estimate the SAT function using the modified Weibull function:

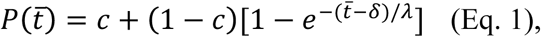

where *P* is the probability of correct response, *c* is a constant reflecting the probability of a correct response at the chance level (*c* was set to 0.5 in our study), 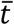 represents the mean response time, and *δ* and λ are free parameters. Here, *δ* captures the point during a trial when *P* rises from chance level, and *λ* captures the slope of the SAT function. We used the method of maximum likelihood to estimate these two parameters. For better convergence, we constrained the parameters such that *δ* must be greater than 0.2 and *λ* must be greater than 0.001.

### Experiment 2: Dynamic combination of sensory and reward information under time pressure

Unlike Experiment 1 in which reward feedback was independent of time, the amount of gain or loss associated with a correct or an incorrect response decreased over time in Experiment 2. In each trial, prior to the onset of color dots, the initial reward value, which was randomly selected from the two possible values (50 and 100 points), was displayed on the screen (Fig. 1B). Therefore, the size of the peripheral arc represented the fraction of this initial reward value over time. Subjects could indicate their decision at any point during a trial.

The reward did not decrease immediately following the onset of dots presentation. Instead, it started to decrease when the estimated SAT rose from the chance level (see the black curve in Figure 7A for an example SAT). This time point was determined for each subject separately based on their SAT and was purposely implemented so that the optimal response time—the peak of the curve in Figure 7B—would not be at the beginning of the dots presentation before subjects could combine sensory and reward information. This ensured that responding as soon as possible was never the optimal decision strategy. It should be noted that subjects were not explicitly informed about the short delay in time that preceded when the reward would start to decrease. However, the subjects received 5 to 10 practice trials after instruction and before the session began, so it is possible they would know that reward did not start to decrease immediately.

The reward value decreased to zero at six different rates independently of the initial reward value. These six decreasing rates were tailored for each subject based on their SAT function estimated from Experiment 1. The subjects were not aware of how decreasing-reward schedules were constructed. Since we were interested in how subjects traded off the accuracy of their judgment against the decreasing reward value, we aimed to set the decrease rates such that the tradeoff was non-trivial. On the one hand, if the rate of decrease was too fast, the subject should respond as soon as possible. On the other hand, if the value decreased too slowly, the subject could comfortably integrate sensory information without losing a significant amount of reward. In both cases, there was no need to actively trade off accuracy with decreasing reward. To avoid these scenarios, we used the optimal model to find the best set of decreasing rates for each subject (see *Optimal model for response time* below). The optimal model predicts, given a subject’s SAT function and a decreasing value function, the optimal response time that maximizes expected gain. For the tradeoff to be non-trivial, the optimal time needs to fall within the range where there is a clear SAT—where the probability of correct response sharply increases. We therefore set the six decreasing rates so that the optimal response time fell within the time window where the probability of correct response was linearly spaced between 0.55 and the subject’s maximum correct probability (e.g., 0.95). For example, if a subject’s maximum correct probability was 0.95, the decreasing rates were set such that the optimal time would correspond to the time points where the probabilities of correct response were 0.55, 0.63, 0.71, 0.79, 0.87, and 0.95.

There were ten blocks of trials, each with 36 trials. Each combination of initial reward value (50 and 100 points) and decreasing rate (six possible rates) was repeated three times in each block while the order of the trials was randomized. Unless otherwise mentioned, trials with both of the two initial reward values were combined for data analysis.

### Optimal model for response time

In Experiment 2, the reward points received after a response at time *t*, *G*(*t,r*), where *r* represents the response outcome (1: correct, 0: incorrect), was positive for correct responses and linearly decreased over time, whereas it was negative for incorrect responses and increased (became less negative) over time:

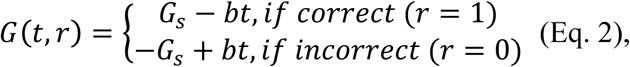

where *G_s_* is the initial reward value, and *b* is the rate at which the reward decreases over time (*b* > 0). For both correct and incorrect response, once the gain or loss reached 0 it stayed at 0.

Due to the SAT, the subject’s performance (probability correct) would improve as a function of time—the more time subjects take, the more sensory information they accumulate over time and the more likely that they would make a correct judgment. Assuming that the subject makes a response at time *t_plan_*, which is a random variable with a mean equal to 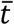, the expected gain at 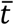 is equal to

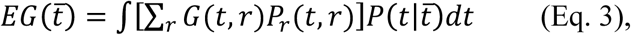

where *r* represents response outcome (1: correct, 0: incorrect), and *P*_*r*_(*t,r*) is the probability of obtaining a response outcome *r* when responding at time *t*. Thus, *P*_*r*_(*t*, 1) is the probability of a correct response at time *t*, whereas *P*_*r*_(*t*, 0) is the probability of an incorrect response is (*P*_*r*_(*t*, 0) =1 − (*P*_*r*_ *t*, 1)). As mentioned above in *Estimating the speed-accuracy tradeoff function*, we used Equation 1 to estimate the probability of a correct response as a function of time. Since *r* is a discrete random variable, marginalizing over *r* at *t* is the sum of all possible values of *r*. On the other hand, 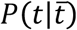 is the probability density function of observing *t* when the mean reaction time is 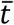. Since *t* is a continuous random variable, marginalizing over *t* is the integral over *t*. An ideal decision maker would choose 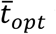 that maximizes the 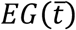 function in Equation 3 (see Fig. 4B).

### Experiment 3. Gambling under time pressure

In Experiment 3, subjects performed a time-dependent gambling task that was mathematically equivalent to the task in Experiment 2, assuming the subjects knew his/her probability of being correct as a function of time. The purpose of this experiment was to provide a control for the dynamic sensory-reward combination studied in Experiment 2 without the need for the subjects to integrate sensory information over time.

Instead of observing color dots presented over time, subjects faced a time-dependent gamble where the probability of winning a monetary reward increased over time (represented by a yellow bar in the center of the screen), while the reward value decreased over time similar to Experiment 2 (Fig. 1C). The probability of winning was tailored for each subject according to his/her SAT function estimated in Experiment 1. This design ensured that the task is mathematically equivalent to the task in Experiment 2 without the need to integrate sensory information over time. The subject could press a key at any point during a trial to accept the gamble with a reward magnitude indicated by the white peripheral arc and reward probability indicated by the yellow central bar. The subjects were not aware of how decreasing-reward schedules were constructed and the fact that the rates of decrease were similar in Experiments 2 and 3. Similar to Experiment 2, reward outcome could be a monetary gain or loss based on the reward magnitude at the time of response. The number of trials and possible combinations were identical to Experiment 2.

### Data analysis

To study the effect of time pressure independently of the integration time, we used RT-matched trials (in terms of mean) from adjacent levels of time pressure. To do so, we first picked all trials from two schedules with adjacent levels of time pressure (e.g., schedules 2 and 3 wherein subjects showed overlapping RTs) for which RTs were between the minimum and maximum RT in the faster and slower schedules, respectively. We then sorted these trials based on their RT and computed the mean RT for each set of trials from the two schedules, and then one by one removed larger RT trials from the slower schedule and smaller RT trials from the faster schedule. This procedure was repeated until the two means of the RTs from the two sets of trials were as close to each other as possible.

To investigate the effect of response time on performance regardless of time windows at the group level (across all subjects), we split response times into ten non-overlapping bins and calculated the mean RT and the proportion of correct trials for each bin (Figs. 2B and 5B). These data were then used to fit performance using a modified Weibull function:

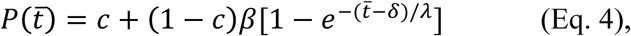

where *P* is the probability of correct response, *c* is a constant reflecting the probability of a correct response at the chance level (*c* was set to 0.5 in our study), 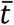 represents the mean response time, and *β*,*δ*, and *λ* are free parameters. Here, *β* captures the steady-state value of *P*, *δ* captures the point during a trial when *P* rises from chance level, and *λ* captures the rising slope of *P*. We used the method of maximum likelihood to estimate these three parameters. For better convergence, we constrained the parameters such that *δ* is greater than 0.2, and *λ* is greater than 0.001.

### Estimating the influence of sample history on choice

To investigate the time-dependent influence of sample dots history on choice on a given trial, we performed a logistic regression analysis on choice (i.e. choose red = 1, green = 0) to estimate the weight of past samples – the proportion of red dots presented in a given time interval (200ms) preceding choice on each trial.

### Comparison of different models for sensory integration under time pressure

To explore neural mechanisms underlying the integration of sensory information in our experiments, we simulated the behavior of various models during sensory integration under time pressure (Experiment 1). More specifically, we simulated the behavior of a model that makes decisions based on stochastic transitions between discrete attractor states (attractor model; Miller & Katz, 2013) in addition to six drift-diffusion models (DDMs).

The attractor model is based on the analysis of coupled groups of neurons in which the decision-making dynamics change from having a single attractor state to a tri-stable system— with the original attractor plus two extra attractors with either one of the groups firing at a high rate—as the input current increases (Wang, 2002). The dynamics then change to a bistable one in which the original attractor has disappeared. To simulate such dynamics with up to three stable attractors, a system of equations using a 6^th^ order difference in firing rates modulated by an urgency signal is needed (Miller & Katz, 2013). Such a system is implemented using the following set of equations:

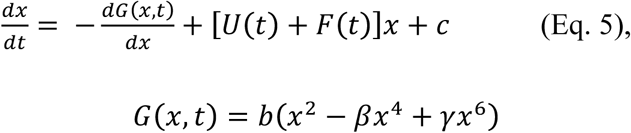

where *x* represents the firing rate, *c* represents white noise (Gaussian noise with mean 0), *b* is the nonlinearity (barrier) parameter that scales all nonlinear terms equally, and *U*(*t*) and *F*(*t)* are the urgency-gating signal and the forcing term, respectively. *U*(*t*) increases linearly from zero upon stimulus onset, and *F*(*t*) is a step function that forces choice before the deadline on a given trial. Decision making in different schedules of Experiment 1 is simulated by using different values of *b* for each time window, corresponding to different levels of time pressure.

In DDMs, the difference between the amounts of evidence supporting the two hypotheses (decision variable) is accumulated until it reaches an upper or lower termination bound (decision boundaries), leading to a choice in favor of one of the two hypotheses. In our experiment, the evidence represents the difference in firing rates between pools of neurons selective for each of the two colors.

We considered six different DDMs. In the basic DDM, the decision variable is updated according to the following equation:

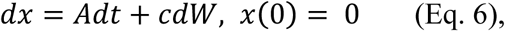

where *x* demonstrates the accumulated value of the difference in evidence, *Adt* represents the constant drift in evidence drawn randomly from a Gaussian distribution with a mean proportional to color strength, and *cdW* represents white noise (Gaussian distribution with mean 0 and variance *c^2^dt*). In this model, different schedules of Experiment 1 are simulated using different decision boundaries for each time window. In the DDM with variant drift rates, we drew drift rate from a Gaussian distribution for each trial.

We also simulated the DDM with collapsing boundaries (Hawkins, Forstmann, et al., 2015). Having collapsing boundaries is equivalent to adding a time-dependent monotonically increasing signal to both signals (evidence) that drive the accumulator. In the DDM with collapsing boundaries, the upper bound *b* at time *t* after onset of evidence accumulation is modeled as:

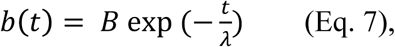

where *λ* is the collapse time constant. Different schedules of Experiment 1 are simulated using variable decision boundaries (*B*) and different time constants of collapse for each time window (*λ*).

We also considered a leak in accumulation of information. The DDM with leak is modeled as in Carland et al. (Carland et al., 2016). To implement the leak in accumulation, we modified the equation for the basic DDM (Eq. 6) to incorporate low-pass filtering of the sensory information using a first-order linear differential equation:

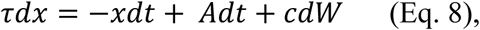

where *τ* is the filter time constant. Similar to the basic DDM, different schedules are simulated using different decision boundaries for each time window.

In another version, the DDM with urgency signal, the decision variable is multiplied by an urgency signal before it is used to make a decision:

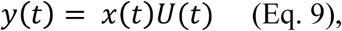

where *U(t)* is the urgency signal that rises linearly from zero with a certain slope. In this model, different schedules are simulated using different decision boundaries for each time window. Finally, the DDM with leak and urgency signal is implemented by incorporating both a leak and an urgency signal into the basic DDM.

We used these models to capture the average SAT function in Experiment 1 (Fig. 2A). The simulations of these six models indicate that an attractor model with stochastic transitions and an urgency signal as well as a DDM with leaky integration and an urgency signal can both qualitatively capture our experimental data (Fig. 4).

### Fitting response time and choice using different DDMs

To determine how the models account for the observed behavior in different schedules, we maximized the likelihood of the choice proportions and the mean and standard deviation of the reaction time from choices (Gaussian error) for each model (Palmer, Huk, & Shadlen, 2005) in 5 bins of reaction time (Fig. 2C). Predictions of each DDM for a given set of parameters were derived from simulating the propagation of the probability density function of the decision variable over time using a simplified one-dimensional Fokker-Planck equation. The attractor model was also fit using the same method as in the DDMs because the attractor model can be estimated by a one-dimensional system of equations with an effective potential (Miller & Katz, 2013).

We note that previous model fitting has been done with purely reaction time data or with data from fixed duration tasks. The task in Experiment 1, however, is a reaction time task but also has a hard deadline. This design makes fitting of choice and reaction data challenging because of the ambiguity in determining the choice if the decision boundary is not crossed before the deadline for a given schedule. We adopted two approaches to tackle this problem and to fit the experimental data. In the first approach, we discarded trials in which the decision boundary was not crossed before the deadline for a given schedule. In the second approach, the sign of the decision variable was used to determine the choice if the decision boundary was not crossed. Fitting of experimental data based on the second approach yielded poor results (data not shown). The main reason for this could be that at none of the presented models could yield very long RTs without low performance. For example, long RTs occur in the DDMs if the drift rate is small which in turn results in low performance. This indicates that additional mechanisms underlie observed long RTs especially for schedules with low levels of time pressure. The results based on the first approach are presented here.

## Acknowledgments

We would like to thank Zohra Aslami, Emily Chu, Brad Duchaine, Clara Guo, and Larry Maloney for helpful comments on the manuscript. This work was supported by NSF EPSCoR (RII Track-2 FEC) grant to AS and by the Ministry of Science and Technology in Taiwan (MOST 101-2628-H-010-001-MY4, 104-2410-H-010-002 -MY3) to SWW.

---

**Supplementary Figure 1.**
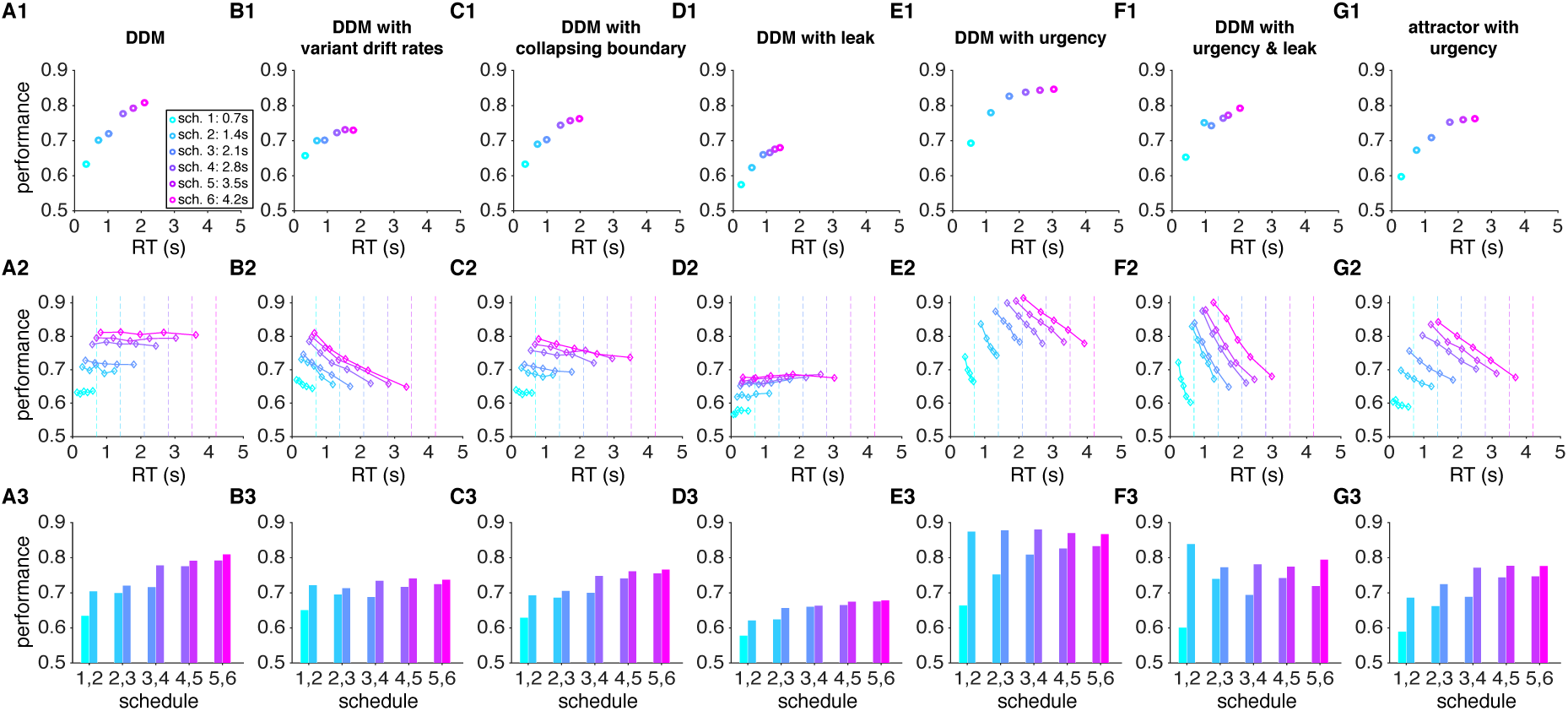
Simulations of experimental data in Experiment 1 using various models. Results for different models are presented on different columns, while panels 1 to 3 of each column plot: the average speed-accuracy tradeoff for a given level of time pressure (panel 1); performance as function of proximity to the deadline (panel 2); and the average performance on RT-matched trials (based on mean) with adjacent levels of time pressure (panel 3). The parameters for each model are selected to show a typical behavior representative of each model. For these simulations, trials in which the decision boundary was not crossed before the deadline were removed from the analysis.

## References

Baird, B., Smallwood, J., Gorgolewski, K. J., & Margulies, D. S. (2013). Medial and lateral networks in anterior prefrontal cortex support metacognitive ability for memory and perception. Journal of Neuroscience, 33(42), 16657–16665.

Battaglia,P. W., & Schrater, P. R. (2007). Humans trade off viewing time and movement duration to improve visuomotor accuracy in a fast reaching task. The Journal of Neuroscience, 27(26), 6984–6994.

Benson, L., & Beach, L. R. (1998). The effects of time constraints on the prechoice screening of decision options. Image Theory: Theoretical and Empirical Foundations, 51–60.

Bogacz, R., Wagenmakers,E. J., Forstmann,B. U., & Nieuwenhuis, S. (2010). The neural basis of the speed–accuracy tradeoff. Trends in Neurosciences, 33(1), 10–16.

Busemeyer,J. R., & Townsend, J. T. (1993). Decision field theory: a dynamic-cognitive approach to decision making in an uncertain environment. Psychological Review, 100(3), 432–459.

Busemeyer,J. R., Wang, Z., & Townsend, J. T. (2006). Quantum dynamics of human decision-making. Journal of Mathematical Psychology, 50(3), 220–241.

Carland,M. A., Marcos, E., Thura, D., & Cisek, P. (2016). Evidence against perfect integration of sensory information during perceptual decision making. Journal of Neurophysiology, 115(2), 915–930.

Chittka, L., Skorupski, P., & Raine, N. E. (2009). Speed–accuracy tradeoffs in animal decision making. Trends in Ecology & Evolution, 24(7), 400–407.

Christopoulos, V., & Schrater, P. R. (2015). Dynamic integration of value information into a common probability currency as a theory for flexible decision making. PLoS Computational Biology, 11(9), e1004402.

Cisek, P., Puskas,G. A., & El-Murr, S. (2009). Decisions in changing conditions: the urgency-gating model. The Journal of Neuroscience, 29(37), 11560–11571.

Dean, M., Wu, S.-W., & Maloney, L. T. (2007). Trading off speed and accuracy in rapid, goal-directed movements. Journal of Vision, 7(5), 1–12.

Deroy, O., Spence, C., & Noppeney, U. (2016). Metacognition in Multisensory Perception. Trends in Cognitive Sciences, 20(10), 736–747.

Ditterich, J. (2006). Stochastic models of decisions about motion direction: behavior and physiology. Neural Networks, 19(8), 981–1012.

Drugowitsch, J., DeAngelis,G. C., Angelaki,D. E., & Pouget, A. (2015). Tuning the speed-accuracy trade-off to maximize reward rate in multisensory decision-making. Elife, 4, e06678.

Durstewitz, D., & Deco, G. (2008). Computational significance of transient dynamics in cortical networks. European Journal of Neuroscience, 27(1), 217–227.

Fleming,S. M., Dolan, R. J., & Frith, C. D. (2012). Metacognition: computation, biology and function. Philosophical Transactions of the Royal Society B: Biological Sciences, 367(1594), 1280–1286.

Fleming,S. M., Huijgen, J., & Dolan, R. J. (2012). Prefrontal contributions to metacognition in perceptual decision making. Journal of Neuroscience, 32(18), 6117–6125.

Fleming,S. M., Weil, R. S., Nagy, Z., Dolan, R. J., & Rees, G. (2010). Relating introspective accuracy to individual differences in brain structure. Science, 329(5998), 1541–1543.

Gao, J., Tortell, R., & McClelland, J. L. (2011). Dynamic integration of reward and stimulus information in perceptual decision-making. PloS One, 6(3), e16749.

Gluth, S., Rieskamp, J., & Büchel, C. (2012). Deciding when to decide: time-variant sequential sampling models explain the emergence of value-based decisions in the human brain. Journal of Neuroscience, 32(31), 10686–10698.

Gold,J. I., & Shadlen, M. N. (2001). Neural computations that underlie decisions about sensory stimuli. Trends in Cognitive Sciences, 5(1), 10–16.

Gold,J. I., & Shadlen, M. N. (2007). The neural basis of decision making. Annual Review of Neuroscience, 30, 535–574.

Griffin, D., & Tversky, A. (1992). The weighing of evidence and the determinants of confidence. Cognitive Psychology, 24(3), 411–435.

Hawkins,G. E., Forstmann, B. U., Wagenmakers, E.-J., Ratcliff, R., & Brown, S. D. (2015). Revisiting the evidence for collapsing boundaries and urgency signals in perceptual decision-making. The Journal of Neuroscience, 35(6), 2476–2484.

Hebart,M. N., Schriever, Y., Donner, T. H., & Haynes, J.-D. (2014). The relationship between perceptual decision variables and confidence in the human brain. Cerebral Cortex, 26(1), 181–130.

Heekeren, H. R., Marrett, S., & Ungerleider, L. G. (2008). The neural systems that mediate human perceptual decision making. Nature Reviews Neuroscience, 9(6), 467–479.

Hernández, A., Zainos, A., & Romo, R. (2002). Temporal evolution of a decision-making process in medial premotor cortex. Neuron, 33(6), 959–972.

Huk,A. C., & Shadlen, M. N. (2005). Neural activity in macaque parietal cortex reflects temporal integration of visual motion signals during perceptual decision making. The Journal of Neuroscience, 25(45), 10420–10436.

Juni,M. Z., Gureckis, T. M., & Maloney, L. T. (2016). Information sampling behavior with explicit sampling costs. Decision, 3(3), 147–168.

Kahneman, D., & Tversky, A. (1979). Prospect Theory: An Analysis of Decision under Risk. Econometrica, 47(2), 263–291.

Katz,L. N., Yates, J. L., Pillow, J. W., & Huk, A. C. (2016). Dissociated functional significance of decision-related activity in the primate dorsal stream. Nature, 535(7611), 285–288.

Kiani, R., Hanks,T. D., & Shadlen, M. N. (2008). Bounded integration in parietal cortex underlies decisions even when viewing duration is dictated by the environment. The Journal of Neuroscience, 28(12), 3017–3029.

Kocher,M. G., & Sutter, M. (2006). Time is money—Time pressure, incentives, and the quality of decision-making. Journal of Economic Behavior & Organization, 61(3), 375–392.

Latimer,K. W., Yates, J. L., Meister, M. L., Huk, A. C., & Pillow, J. W. (2015). Single-trial spike trains in parietal cortex reveal discrete steps during decision-making. Science, 349(6244), 184–187.

Luce, R. D. (1986). Response times: Their role in inferring elementary mental organization. Oxford University Press.

Maule,A. J., Hockey, G. R. J., & Bdzola, L. (2000). Effects of time-pressure on decision-making under uncertainty: changes in affective state and information processing strategy. Acta Psychologica, 104(3), 283–301.

Miller, P., & Katz, D. B. (2010). Stochastic transitions between neural states in taste processing and decision-making. The Journal of Neuroscience, 30(7), 2559–2570.

Miller, P., & Katz, D. B. (2013). Accuracy and response-time distributions for decision-making: Linear perfect integrators versus nonlinear attractor-based neural circuits. Journal of Computational Neuroscience, 35(3), 261–294.

Ordonez, L., & Benson, L. (1997). Decisions under time pressure: How time constraint affects risky decision making. Organizational Behavior and Human Decision Processes, 71(2), 121–140.

Palmer, J., Huk,A. C., & Shadlen, M. N. (2005). The effect of stimulus strength on the speed and accuracy of a perceptual decision. Journal of Vision, 5(5), 376–404. https://doi.org/10.1167/5.5.1

Payne,J. W., Bettman, J. R., & Johnson, E. J. (1993). The adaptive decision maker. Cambridge University Press.

Ratcliff, R. (1978). A theory of memory retrieval. Psychological Review, 85(2), 59–108.

Ratcliff, R., & McKoon, G. (2008). The diffusion decision model: theory and data for two-choice decision tasks. Neural Computation, 20(4), 873–922.

Ratcliff, R., & Rouder, J. N. (1998). Modeling response times for two-choice decisions. Psychological Science, 9(5), 347–356.

Ratcliff, R., & Smith, P. L. (2004). A comparison of sequential sampling models for two-choice reaction time. Psychological Review, 111(2), 333.

Rieskamp, J., & Hoffrage, U. (2008). Inferences under time pressure: How opportunity costs affect strategy selection. Acta Psychologica, 127(2), 258–276.

Roitman,J. D., & Shadlen, M. N. (2002). Response of neurons in the lateral intraparietal area during a combined visual discrimination reaction time task. The Journal of Neuroscience, 22(21), 9475–9489.

Shadlen,M. N., & Newsome, W. T. (2001). Neural basis of a perceptual decision in the parietal cortex (area LIP) of the rhesus monkey. Journal of Neurophysiology, 86(4), 1916–1936.

Stanford,T. R., Shankar, S., Massoglia, D. P., Costello,M. G., & Salinas, E. (2010). Perceptual decision making in less than 30 milliseconds. Nature Neuroscience, 13(3), 379–385.

Thura, D., Beauregard-Racine, J., Fradet, C.-W., & Cisek, P. (2012). Decision making by urgency gating: theory and experimental support. Journal of Neurophysiology, 108(11), 2912–2930.

Thura, D., Cos, I., Trung, J., & Cisek, P. (2014). Context-dependent urgency influences speed–accuracy trade-offs in decision-making and movement execution. The Journal of Neuroscience, 34(49), 16442–16454.

Tsetsos, K., Usher, M., & McClelland, J. L. (2011). Testing multi-alternative decision models with non-stationary evidence. Frontiers in Neuroscience, 5, 63.

Usher, M., & McClelland, J. L. (2001). The time course of perceptual choice: the leaky, competing accumulator model. Psychological Review, 108(3), 550–592.

Wang, X.-J. (2002). Probabilistic decision making by slow reverberation in cortical circuits. Neuron, 36, 955–968.

Wu, S.-W., Delgado,M. R., & Maloney, L. T. (2015). Gambling on visual performance: neural correlates of metacognitive choice between visual lotteries. Frontiers in Neuroscience, 9, 314.

Yates,J. L., Park, I. M., Katz, L. N., Pillow, J. W., & Huk, A. C. (2017). Functional dissection of signal and noise in MT and LIP during decision-making. Nature Neuroscience, 20(9), 1285–1292.

